# EGR1 Mediates Riluzole-Induced Apoptosis in Osteosarcoma via the Yap/p73-Bax Signaling Axis

**DOI:** 10.1101/2025.10.15.682608

**Authors:** Syeda Maryam Azeem, Shraddha ChandThakuri, Pooja Prakash Rao, Tamia Abbasi, Jannae McFarlane, Raisa Munira, Sahanama Porshe, Vinagolu K. Rajasekhar, John H. Healey, Shahana S. Mahajan

**Author notes:** Corresponding Authors: Shahana S. Mahajan.

## Abstract

Osteosarcoma (OS), although rare, is the most common primary bone cancer, primarily affecting individuals aged 10-30 years. Despite therapeutic advances, survival rates have remained stagnant for decades. Recent studies show that Riluzole, a glutamate receptor antagonist, induces apoptosis in OS cells both *in vitro* and *in vivo*. Our previous work demonstrated that Riluzole increases reactive oxygen species (ROS), activating c-Abl kinase, which phosphorylates Yes-associated protein (Yap) at tyrosine 357. This modification promotes nuclear translocation of Yap and interaction with p73, enhancing Bax expression and inducing apoptosis. Early Growth Response 1 (*EGR1*), a zinc finger transcription factor often linked to apoptosis in other cancers, is significantly downregulated in OS. Here, we investigated the role of *EGR1* in Riluzole-mediated apoptosis across OS cell lines and patient-derived xenografts (PDX). In this study, we show that Riluzole upregulates *EGR1* expression in all OS cell lines. Chromatin immunoprecipitation followed by qPCR confirmed that *EGR1* directly binds to the Bax promoter along with Yap/p73, enhancing Bax expression. Immunohistochemistry of *in vivo* xenograft tumors from Riluzole-treated mice revealed increased *EGR1* and cleaved caspase-3 levels, indicating elevated apoptosis, while reduced NUMA expression suggested diminished tumor proliferation. Together, these findings reveal a novel mechanism where Riluzole promotes apoptosis through upregulation of *EGR1*, which then cooperates with YAP/p73 to activate Bax expression. These insights establish Riluzole as a promising therapeutic intervention for OS treatment through modulation of the *EGR1*/Yap/p73/Bax signaling axis.

## BACKGROUND

Osteosarcoma (OS), while a rare cancer, is the most common primary bone malignancy in adolescents and young adults, with the highest incidence occurring between the ages of 10 and 14 (1). Tumors are frequently seen localized in the long tubular bones during the growth phase, mostly seen in the knee joint and can also occur in the pelvic bone area (2). Localized tumors can metastasize to other organs even before the patient is diagnosed but 90% of the time, OS metastasizes to lungs (2). Unfortunately, very few diagnostic/prognostic biomarkers are available for proper early detection or treatment response for OS. Less than 1000 cases are reported per year in the US and overall incidence ranges anywhere between 4.4-5.6 per million per year worldwide (3). Primary tumor resection and neoadjuvant chemotherapy for improved overall survival rate of OS is still the mainstay treatment for three decades (4). Concurrently, the chemotherapeutic treatment regime consists of a combination of drugs such as doxorubicin, methotrexate, and cisplatin after tumor resection (4, 5). Even with a 5-year survival rate, which is ∼70% for localized tumors, the already metastasized tumors still remain the biggest concern (3). With the failing conventional therapies and the multi-drug resistance emergence, there is an urgent need to develop targeted drug therapies, prognostic and diagnostic biomarkers to increase the overall survival rate of OS. Lately, the FDA-approved drug Riluzole, used for ALS, is being investigated for its ability to induce apoptosis in many cancer types (6, 7). Our lab investigated Riluzole’s potential for OS therapy.

Riluzole is used for the treatment of Amyotrophic Lateral Sclerosis (ALS) and it acts by inhibiting glutamate release (8). Physiologically, glutamate, the major excitatory neurotransmitter, is involved in several neural and metabolic processes (9). In bone cells, glutamate is important during bone cell differentiation and development (10). The family of glutamate receptors are involved in abnormal signaling in various cancers such as melanoma, gliomas and OS (11-13). Previous studies have shown that inhibiting glutamate release causes a significant decrease in the cancer progression and migration (14, 15). Riluzole has been shown to inhibit glutamate release in several cancer cell types (6, 14, 16). It also decreases cell proliferation and survival leading to apoptosis in cell culture and *in vivo* mouse studies for OS, melanomas and gliomas (6, 16). Although proven to be effective in various cancer cells, the mechanism of Riluzole is still being investigated. Recently, we showed that Riluzole induces apoptosis in LM7 metastatic OS cells (17). The study found that Riluzole treatment increases cell stress generating reactive oxygen species (ROS) which activates c-Abl kinase that in-turn phosphorylates Yap at Y357 residue. The phosphorylated Yap along with p73 binds to the Bax-promoter region and increases Bax protein expression leading to apoptotic cell death in OS (17).

Early growth response 1 (*EGR1*), a zinc finger transcription factor, is a 543 amino acid long protein and 57kDa in size that binds to GC rich regions on the promoter region of their target genes (18). *EGR1* gene is highly conserved in humans and is regulated by external stimuli such as cell stress, hypoxia, and UV-radiation (18, 19). *EGR1* gene is involved in the regulation of differentiation, proliferation, apoptosis and has been shown to have tumor suppressor activity in several carcinoma cells except prostate cancer (19-22). *EGR1* has recently been indicated to play a prognostic role in breast cancer (23). In prostate cancer, *EGR1* is highly expressed in tumor cells indicating its involvement in tumor progression (19, 21). However, Zagurovskaya et al. has shown that over-expression of *EGR1* in prostate cancer sensitizes cells to radiation induced DNA damage that leads to binding of *EGR1* to the Bax-promoter along with Yap and increasing Bax expression (24). In most cancer types, the *EGR1* interaction with Bax-promoter was shown to be necessary to cause apoptosis (22). This led us to question if *EGR1* is involved in our previously described mechanism of Riluzole-induced apoptosis in OS. In OS and other cancers, *EGR1* gene is highly downregulated, and its upregulation causes increased apoptosis (22, 25). In other cancer types and OS, increase in cell stress caused increased generation of reactive oxygen species (ROS) which in turn increased *EGR1* expression leading to apoptosis (26, 27). Interestingly, Stuart et. al. 2005, has shown that c-Abl regulates *EGR1* through ROS in U2OS cells. We have previously shown in OS that Riluzole induces apoptosis via the activation of c-Abl kinase (17). However, the role of *EGR1* in Riluzole-induced apoptosis has not been demonstrated. Several studies have shown interaction of transcription factor *EGR1* with Bax-promoter region along with Yap in different cancer cells that leads to apoptosis except in OS (22, 24). Interestingly, *EGR1* expression increased in different OS cells causing apoptosis upon treatment with chemotherapeutic agents (20). However, Riluzole-induced *EGR1* expression in the apoptosis of OS is still unexplored leading us to hypothesize that *EGR1* may play a role in Riluzole induced apoptosis in OS. Here we present evidence that Riluzole treated OS cell lines and OS patient derived xenograft (PDX) lines show a significant increase in *EGR1* expression that leads to increased binding of *EGR1* to the Bax-promoter region which in turn facilitates Yap/73 binding and increased Bax expression, ultimately leading to apoptosis in osteosarcoma.

## RESULTS

### Riluzole transcriptionally upregulates pro-apoptotic genes and suppresses pro-proliferative genes in OS

Our previous research established that Riluzole promotes the expression of Yap and p73, and that their recruitment to, pro-apoptotic gene, the Bax promoter region enhances Bax transcription, leading to apoptosis in LM7 cells (17). Interestingly, in prostate cancer, it has been shown that Yap interacts with *EGR1* through its WW domain, playing a role in Bax-mediated apoptosis (24). Building on these findings, we sought to investigate whether *EGR1* is similarly involved in the Yap/p73 complex in the context of Riluzole-induced apoptosis in OS. To explore this possibility, we utilized two human osteosarcoma cell lines: U2OS (Figure 1A), which represents a primary tumor and expresses wild-type p53, and Saos-2 derived LM7 (Figure 1B), a lung metastatic derivative that lacks functional p53. To assess the involvement of *EGR1*, we first treated both the above cell lines with Riluzole and measured *EGR1* mRNA expression. In U2OS cells (Figure 1A), we observed a robust induction of *EGR1* expression within one hour of Riluzole treatment. In contrast, LM7 cells (Figure 1B) showed a delayed but significant increase in *EGR1* expression, peaking at four hours post Riluzole treatment. These findings indicate that *EGR1* is responsive to Riluzole treatment in both primary and metastatic OS models.

**Figure 1.**
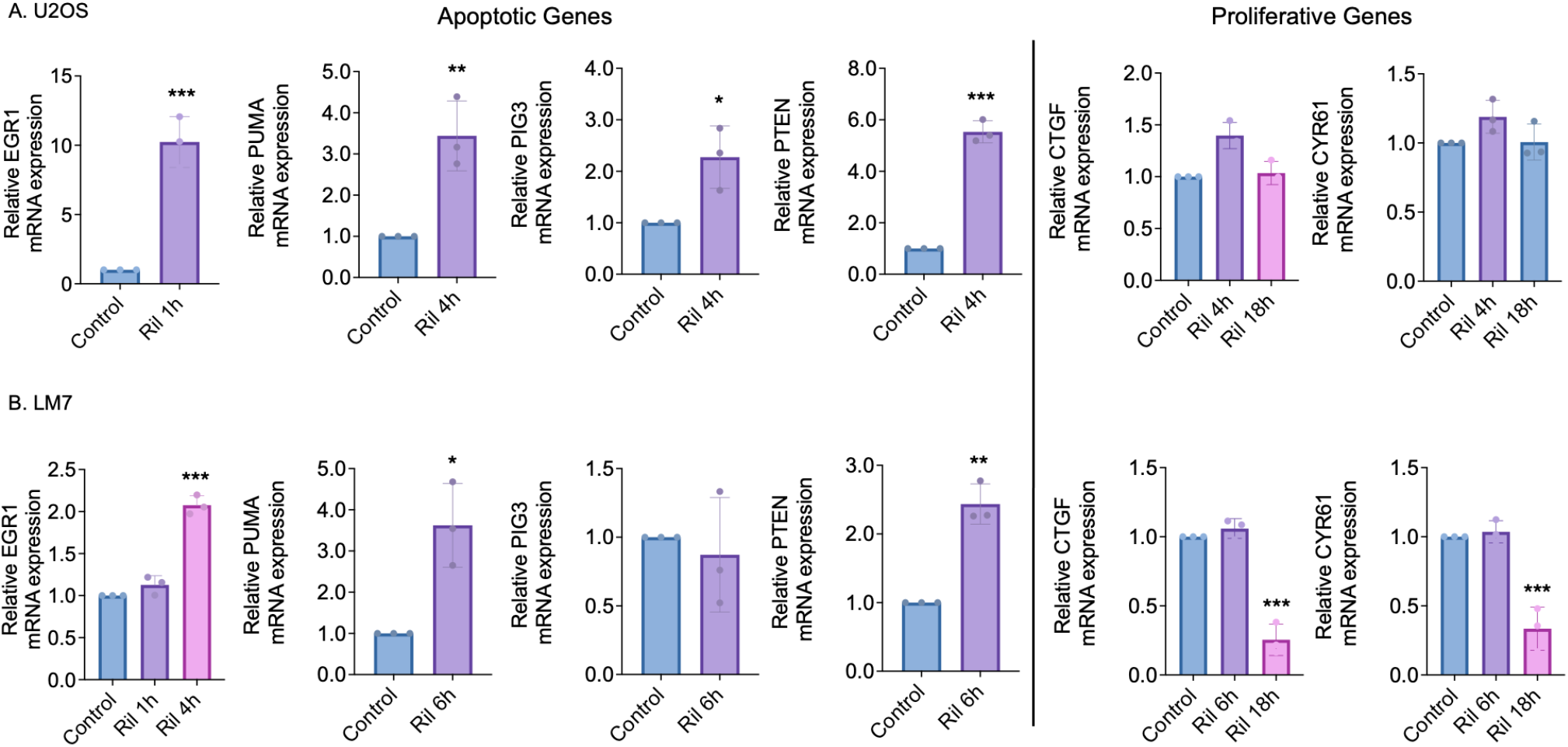
Transcriptional upregulation of apoptotic genes and downregulation of proliferative genes by Riluzole in OS: Quantitative RT-PCR analysis showing relative mRNA expression levels of apoptotic genes: *EGR1, PUMA, PIG3*, and *PTEN* and proliferative genes: *CYR61* and *CTGF* in **(A)** U2OS and **(B)** LM7 cells following 100 µM Riluzole treatment for the indicated time points. The values were calculated using three different biological sample data and the p-values were calculated using two-tailed Student’s t-test (*p<0.05, **p<0.002, ***p<0.001).

Given our earlier observation that Bax expression is elevated following Riluzole treatment in LM7 cells, we next examined whether other pro-apoptotic genes regulated by *EGR1* might also be upregulated. Specifically, we assessed the expression of *PUMA, PIG3*, and *PTEN* genes known to be transcriptionally regulated by *EGR1* (28, 29). All three genes exhibited increased expression in U2OS cells following Riluzole treatment, supporting the hypothesis that *EGR1* contributes to a broader apoptotic gene expression in OS. Interestingly, in LM7 cells, we saw increase only in *PUMA* and *PTEN* expression but not *PIG3* expression. In addition to the pro-apoptotic genes, we evaluated the expression of pro-proliferative genes such as *CYR61* and *CTGF*, which are often implicated in tumor progression and are well known downstream targets of the Yap/TEAD binding complex (30-32). In U2OS cells, we did not see any increase in the expression of *CYR61* or *CTGF*. In LM7 cells, both *CYR61* and *CTGF* were significantly downregulated in response to Riluzole treatment. This suggests a shift in the cellular program from proliferation to apoptosis, potentially mediated by *EGR1* activity.

### Riluzole treatment increases EGR1 expression in OS cell lines

Previously it was reported that *EGR1* is downregulated in human OS cells, *in vitro*, and treatment with traditional chemotherapeutic drugs such as Doxorubicin, Methotrexate and Cisplatin increased *EGR1* expression causing apoptosis (20). We hypothesized that Riluzole treatment may similarly enhance *EGR1* expression, given its ability to induce apoptosis in OS cells. To validate this, we examined the basal expression levels of *EGR1* across various OS cell lines including metastatic and non-metastatic cell lines (Figure 2A). Our results revealed that *EGR1* mRNA basal expression was significantly lower in all OS cell lines compared to the normal human fetal osteoblast (hFOB) cell line. At the protein level, however, *EGR1* expression was not suppressed in Saos-2, HOS, and 143B cell lines. In contrast, LM7, U2OS, MG63, and MG63.3 showed significantly lower *EGR1* protein expression compared to hFOB (Figure 2A).

**Figure 2.**
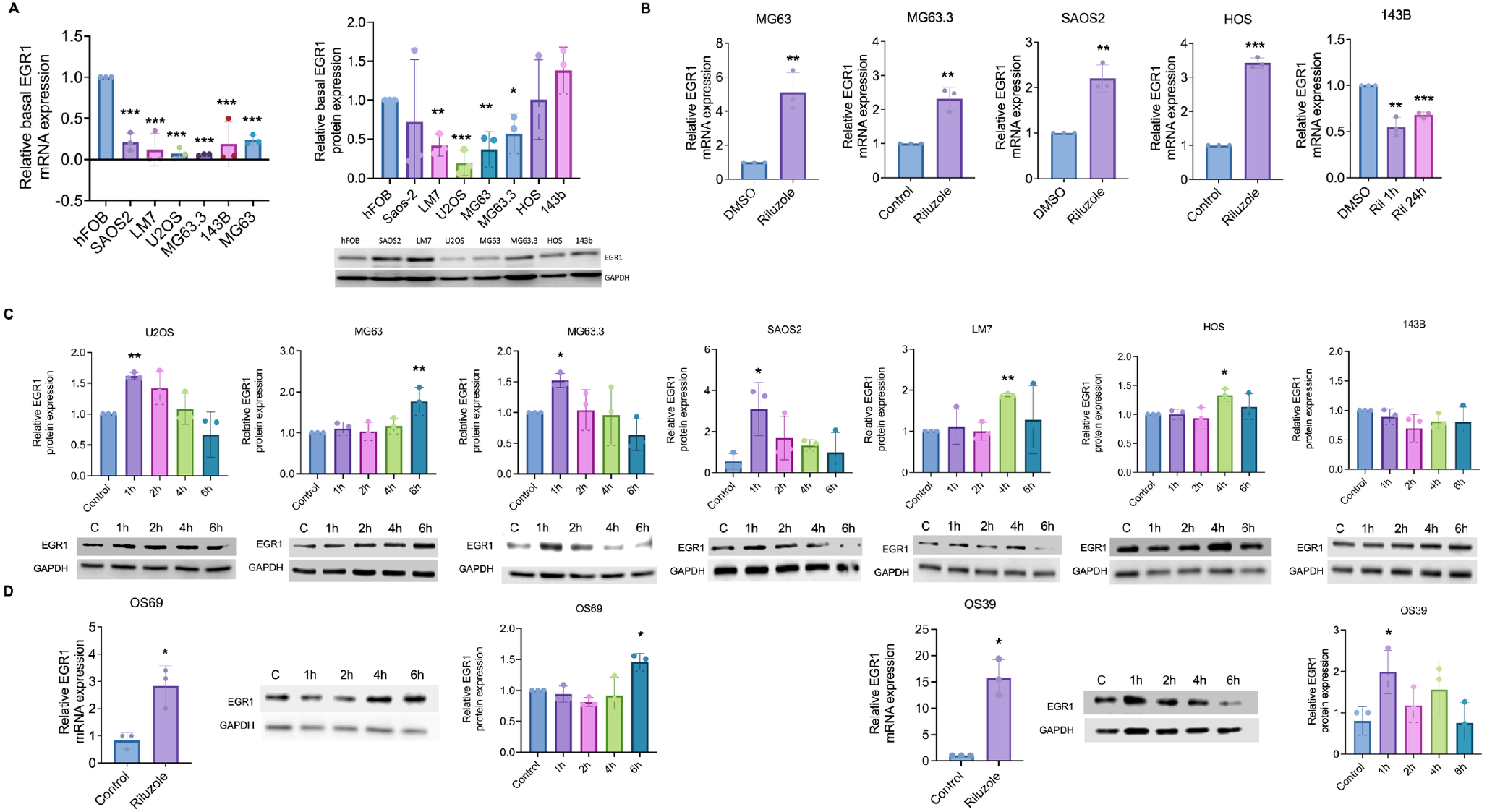
Regulation of EGR1 expression in osteosarcoma by Riluzole: **(A)** The bar graph shows the relative mRNA and protein levels of *EGR1* across untreated osteosarcoma cell lines compared to hFOB (control). Data is expressed as fold change relative to hFOB. **(B)** Quantitative RT-PCR analysis showing *EGR1* expression in multiple osteosarcoma cell lines after 1 hour of 100 µM Riluzole treatment. Data is expressed as fold change relative to control. **(C)** Representative Western blot analysis and corresponding quantification of *EGR1*/GAPDH ratios after 100 µM Riluzole treatment at the indicated time points in different osteosarcoma cell lines. **(D)** The bar graph shows the relative mRNA levels of *EGR1* (left panel) across two PDX lines after 1 hour of 100 µM Riluzole treatment and relative protein *EGR1* expression (right panel) after 100 µM Riluzole treatment at indicated time points. Data is expressed as fold change relative to control from three independent experiments and were analyzed for statistical significance using two-tailed Student’s t-test and one-way ANOVA with Bonferroni’s post-hoc analysis (*p<0.03, **p<0.002, ***p<0.0001).

We then investigated the effect of Riluzole on *EGR1* expression and at the mRNA level (Figure 2B), we observed a significant increase in *EGR1* expression across a series of OS cell lines after 1 hour of treatment, except for 143B. At the protein level (Figure 2C), *EGR1* expression significantly increased after 1 hour of treatment in MG63.3, U2OS, and Saos-2 cells. In LM7 and HOS cells, a significant increase in *EGR1* expression was observed after 4 hours of treatment, whereas MG63 cells exhibited a significant increase after 6 hours. Interestingly, the 143B cell line showed no significant change in *EGR1* expression at either the mRNA or protein level, even after 24 hours of Riluzole treatment. To further explore our hypothesis, we examined the effects of Riluzole on *EGR1* expression in two PDX lines: OS69 and OS39 (Figure 2D). Our results showed a 3-fold increase in *EGR1* mRNA expression within 1 hour of Riluzole treatment and a nearly 2-fold increase in protein expression after 6 hours of treatment in OS69 PDX line. On the other hand, in OS39 PDX line, we observed 15-fold increase in *EGR1* mRNA expression and 2-fold increase in protein expression within 1 hour of treatment. These findings suggest that Riluzole treatment enhances *EGR1* expression in OS cells as well as PDX lines, albeit at an accepted levels of variations in the onset of *EGR1* expression across different cell lines.

### Riluzole-induced EGR1 expression through ROS generation

Previous studies have shown that ROS sensitizes increase in *EGR1* expression (26, 27). In addition, it was shown that increase in *EGR1* could partly be mediated by ROS activated c-Abl kinase in U2OS (33). On the other hand, our recent study demonstrated that ROS production increases upon Riluzole treatment in both LM7 and U2OS (17, 34). Therefore, we sought to determine whether ROS enhances *EGR1* expression and whether Riluzole-induced ROS directly stimulates *EGR1* expression. We examined *EGR1* levels in samples treated with Riluzole alone, NAC alone, and a combination of Riluzole and NAC. We showed that there was a significant increase in *EGR1* expression following Riluzole treatment in both U2OS and LM7 cells (Figure 3). However, co-treatment with Riluzole and the antioxidant NAC led to a marked reduction in *EGR1* expression compared to Riluzole alone, indicating a counteractive effect. These findings confirm that upon Riluzole treatment, ROS directly stimulates *EGR1* expression.

**Figure 3.**
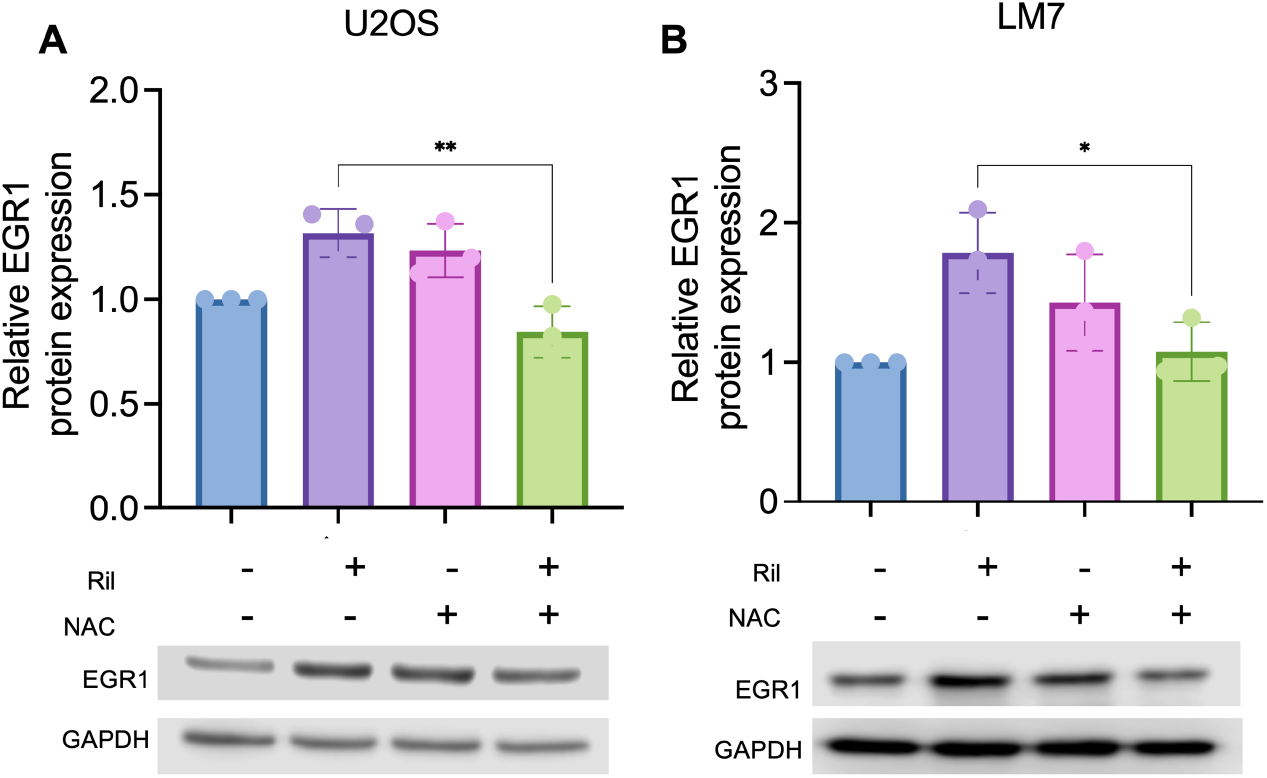
Riluzole-induced EGR1 expression is mediated by ROS generation in OS: Quantification of *EGR1* protein levels in, **(A) U2OS** and **(B) LM7** cells treated with control, Riluzole, NAC, and Riluzole combined with NAC. The bar graph represents the *EGR1*/GAPDH band intensity ratio from three independent experiments. Representative Western blot images below each graph show *EGR1* and GAPDH bands for each treatment group. The values were calculated using three different biological sample data and the p-values were calculated using one-way ANOVA with Bonferroni’s post-hoc analysis (*p<0.03, **p<0.002, ***p<0.0001)

### EGR1 facilitates the binding of Yap/p73 on the Bax promoter to increase its expression upon Riluzole treatment

Zakurovskaya et al. (2009) showed that exogenous overexpression of *EGR1* sensitizes prostate cancer cells to radiation, promoting Yap/*EGR1* binding to the Bax promoter inducing Bax-driven apoptosis. In our earlier work (17), we demonstrated that Riluzole induces apoptosis in OS by increasing Bax expression through the binding of Yap/p73 to the Bax promoter. However, Yap or p73 may not bind to the DNA directly in a sequence specific manner. This prompted us to investigate whether the Yap/p73 complex binding to the Bax promoter is mediated by *EGR1* in OS cells treated with Riluzole. In pursuit of this, we conducted chromatin immunoprecipitation followed by qPCR (ChIP-qPCR) using Bax-promoter primers. This approach allowed us to analyze the direct, endogenous binding of Riluzole-induced *EGR1* to the Bax promoter by complexing with Yap and p73. Additionally, we used pol II antibodies to pull down DNA fragments bound to pol II, ensuring that the immunoprecipitated fragments were in an actively transcribing state (Figure 4A,B).

**Figure 4.**
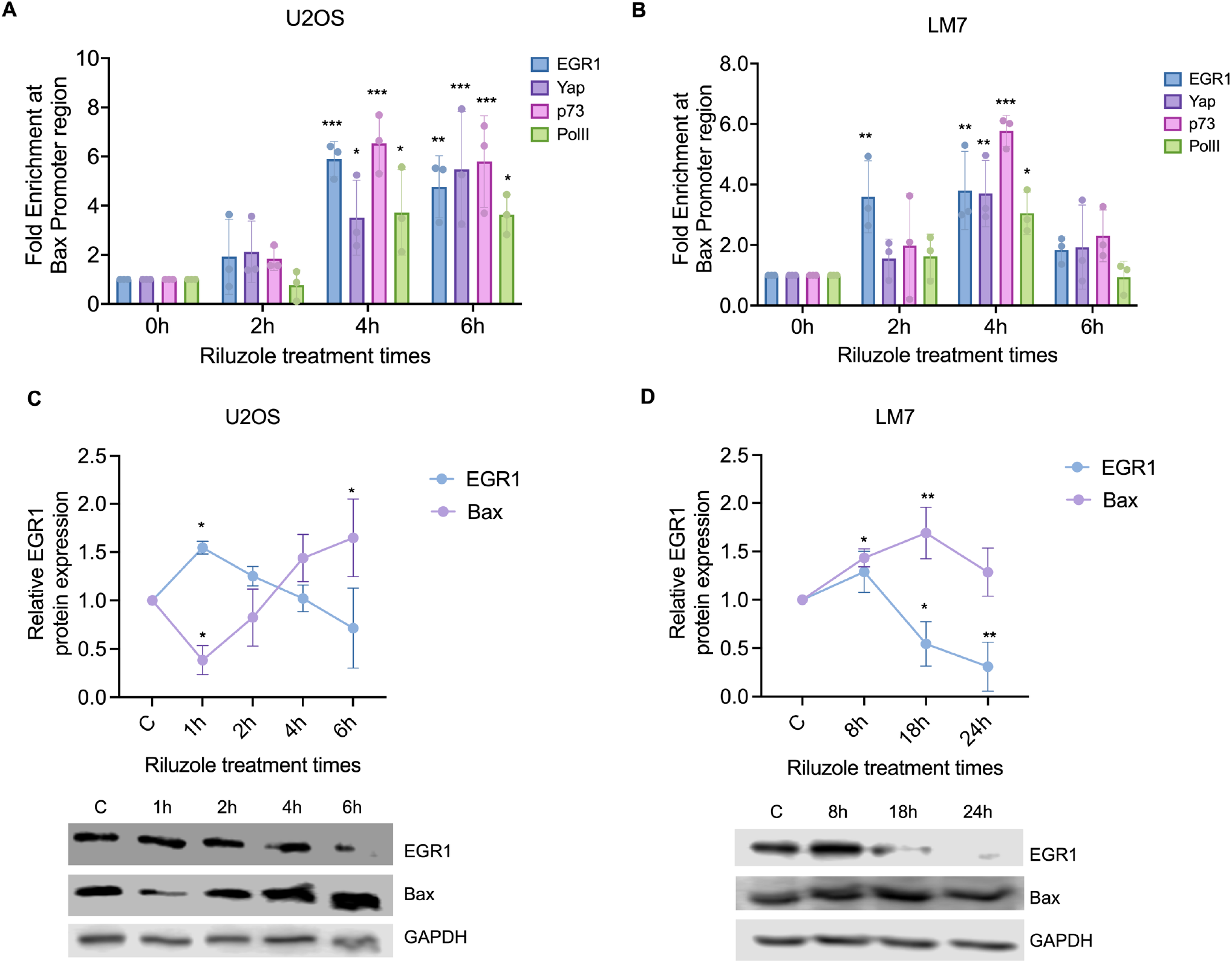
Riluzole upregulates Bax expression via EGR1/Yap/p73 axis in OS. Chromatin immunoprecipitation (ChIP) qPCR analysis showing the fold enrichment of *EGR1*, Yap, p73, and Pol II binding at the Bax promoter region in **(A)** U2OS and **(B)** LM7 cells at 0, 2, 4, and 6 hours after Riluzole treatment. Western blot and densitometry analysis of *EGR1* and Bax protein expression levels in **(C)** U2OS and **(D)** LM7 cells following Riluzole treatment for the indicated times. GAPDH was used as a loading control. The bar graphs represent the *EGR1*/GAPDH and Bax/GAPDH ratios, showing a time-dependent increase in protein expression levels. The values were calculated using three different biological sample data and the p-values were calculated using two-way ANOVA (*p < 0.03, **p < 0.002, ***p<0.0002).

Previously, we observed a significant increase in the fold enrichment of the Yap/p73 complex on the Bax promoter region (17). Notably, in our current study, in U2OS, Riluzole treatment led to a 4-to 6-fold enrichment of the *EGR1*/Yap/p73 complex on the Bax promoter after 4 and 6 hours of treatment (Figure 4A). Similarly, in LM7, a 4- to 6-fold enrichment of the *EGR1*/Yap/p73 complex on the Bax promoter was observed after 4 hours of Riluzole treatment (Figure 4B). These results confirm that the *EGR1* is the key mediator facilitating Yap/p73 binding on the Bax promoter, enhancing its expression and driving Riluzole-induced apoptosis in OS.

Our ChIP assay data revealed that *EGR1* binds to Bax promoter along with Yap/p73 following Riluzole treatment. To investigate whether this resulted in an increase in Bax expression, we re-probed the same blots with a Bax antibody to confirm *EGR1* expression. An increase in Bax expression was indeed observed towards the end of the 6-hour Riluzole treatment. Notably, *EGR1* expression increased within the first hour of treatment and decreased by the 6-hour mark in U2OS (Figure 4C). In contrast, Bax expression began to increase within 6 hours of treatment.

In metastatic LM7 (data not shown), no early increase in Bax expression was observed that remains consistent with delayed rise in *EGR1* protein levels, which started after 4 hours of Riluzole treatment. Our previous study (17) indicated that Bax expression increases after 24 hours of treatment. To determine the starting point of this increase, LM7 cells were treated with Riluzole for 8, 18, and 24 hours (Figure 4D). Bax expression showed a significant increase after 8 hours of treatment, peaking at 18 hours. This temporal pattern suggests that the Riluzole-induced activation of *EGR1* and its subsequent binding to the Bax promoter contribute to the upregulation of Bax expression, which is associated with the pro-apoptotic effects observed in osteosarcoma cells.

### Increase in Riluzole induced EGR1 expression in *in vivo* mouse models

While the above *in vitro* findings demonstrate that Riluzole induces *EGR1* expression and promotes apoptosis in above osteosarcoma cells, we extended our study into *in vivo* mouse models for establishing physiological relevance and translational potential of this drug. Riluzole is metabolized in the liver via the cytochrome P450 1A2 (CYP1A2) enzyme (35). The heterogenous expression of CYP1A2 enzyme across the human population contributes to resulting variability in therapeutic response to Riluzole and its rapid metabolism in humans present limitations for its clinical application (35). Therefore, we included two Riluzole prodrugs (36), FC3423 and TroRiluzole, engineered for improved stability and bioavailability, in our *in vivo* studies to assess whether they also induce *EGR1* expression and apoptotic signaling. Comparing *EGR1* expression profiles and therapeutic outcomes across Riluzole and its prodrugs in xenograft mouse models may allow us to evaluate not only the mechanistic role of *EGR1* in tumor suppression in general but also the relative efficacy of these compounds as viable therapeutic agents for osteosarcoma.

We performed MTT assay to determine the effect of Riluzole and its pro-drugs, FC3423 and TroRiluzole, in LM7 *in vitro* (Figure 5A). We observed that the apoptotic rate of cells reached 50% at 67 µM for Riluzole, 32 µM for FC3423 and 87 µM for TroRiluzole (Figure 5B). LM7 cells were more sensitive to FC3423 than Riluzole and TroRiluzole, *in vitro*. We sought to confirm our *in vitro* data by using OS mouse models. Tail vein xenografts were established in 5-week-old female mice (n=5) using LM7 cells. Subsequently, mice were randomly placed into 4 groups after the development of tumors: Control(G1), Riluzole(G2), FC3423(G3), TroRiluzole(G4). Drugs were administered through oral gavage daily to maintain steady levels of drug in the blood plasma. Tumor volume was measured every week for all the groups, and the bioluminescence data was recorded for both ventral and dorsal flux. All animals were sacrificed at the end of Week 4 due to tumor burden in the control group. Our results demonstrated that all the treatment groups saw a decrease in the total bioluminescence flux compared to the control group. Interestingly, TroRiluzole treatment resulted in a significant decrease in the total bioluminescence flux compared to Riluzole or FC3423 treated group at the end of Week 4 (Figure 5C, D).

**Figure 5.**
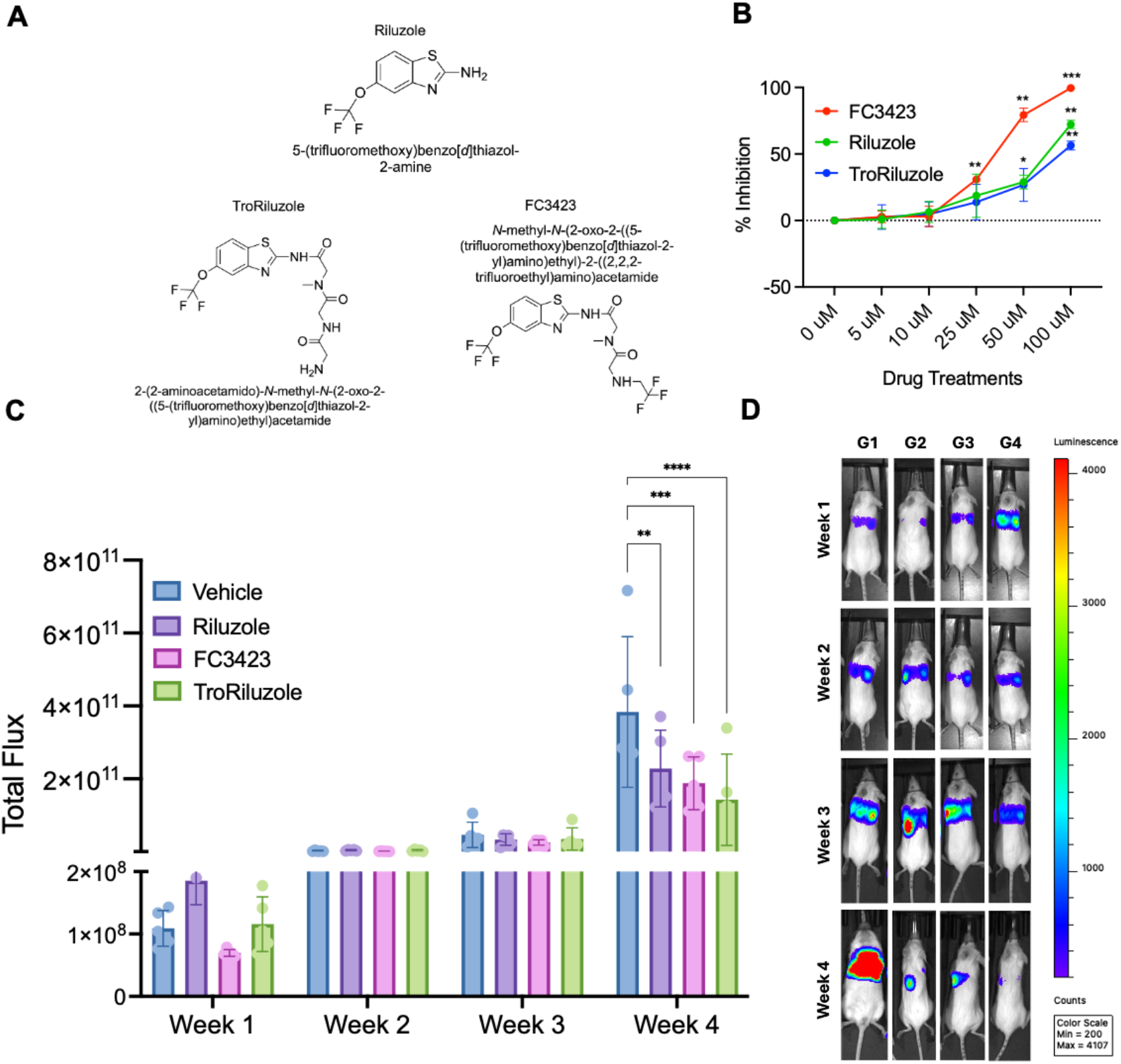
Riluzole and its pro-drugs inhibit the LM7 cell line viability *in vitro* and reduce tumor progression *in vivo*. **(A)** Chemical structures of Riluzole, TroRiluzole, and FC3423 administered for investigating their effectiveness on *in vitro* cell viability and on *in vivo* metastases of osteosarcoma. **(B)** Cytotoxicity assay: LM7 cells were treated with different indicated concentrations of Riluzole, TroRiluzole and FC3423. After 48 hours, the cells were incubated for an hour with MTT reagent, and the precipitate was dissolved with DMSO. The plate was read, and the data was plotted. Data are represented as mean SD of three biological replicates, with statistical analysis performed using one-way ANOVA with Dunnett’s post-hoc analysis (*p<0.03, **p<0.002, ***p<0.0001) **(C)** Bioluminescence data of animals plotted as change in the total flux each week. Data are represented as mean SD of n=5/group, with statistical analysis performed using two-way ANOVA with Dunnett’s post-hoc analysis (*p<0.03, **p<0.002, ***p<0.0001). **(D)** Representative mice images of each group showing dorsal flux among the different groups from week 1 to week 4.

To validate our *in vitro* findings across various osteosarcoma (OS) cell lines, we then examined the *in vivo* effects of Riluzole and its prodrugs, FC3423 and TroRiluzole, on tumor tissues from xenografted mice with pulmonary metastases. Immunohistochemical analyses were conducted to assess markers of cell proliferation and apoptosis, as well as the expression of *EGR1*. Nuclear mitotic apparatus protein (NUMA) staining revealed a significant reduction in cell proliferation in all drug-treated groups compared to controls (Figure 6A). Additionally, cleaved caspase-3 staining indicated a marked increase in apoptosis within the tumor tissues of treated mice (Figure 6B). These results suggest that Riluzole and its prodrugs effectively suppress tumor growth by inhibiting proliferation and promoting apoptotic pathways. These findings are also consistent with our previous study indicating that Riluzole effectively induces apoptosis and reduces tumor size in osteosarcoma models (37). Furthermore, *EGR1* expression was notably elevated in the treated groups, with a three-fold increase observed in the Riluzole-treated mice, a 4-8-fold increase in the FC3423 group, and a 3-6-fold increase in the TroRiluzole group (Figure 6C). This *in vivo* upregulation of *EGR1* is consistent with our *in vitro* findings, reinforcing its potential role in mediating diminished tumor growth and increased apoptosis in OS xenograft tumor tissues. Together, all the above data supports the therapeutic potential of Riluzole and the importance of upregulating *EGR1* in OS treatment strategies.

**Figure 6.**
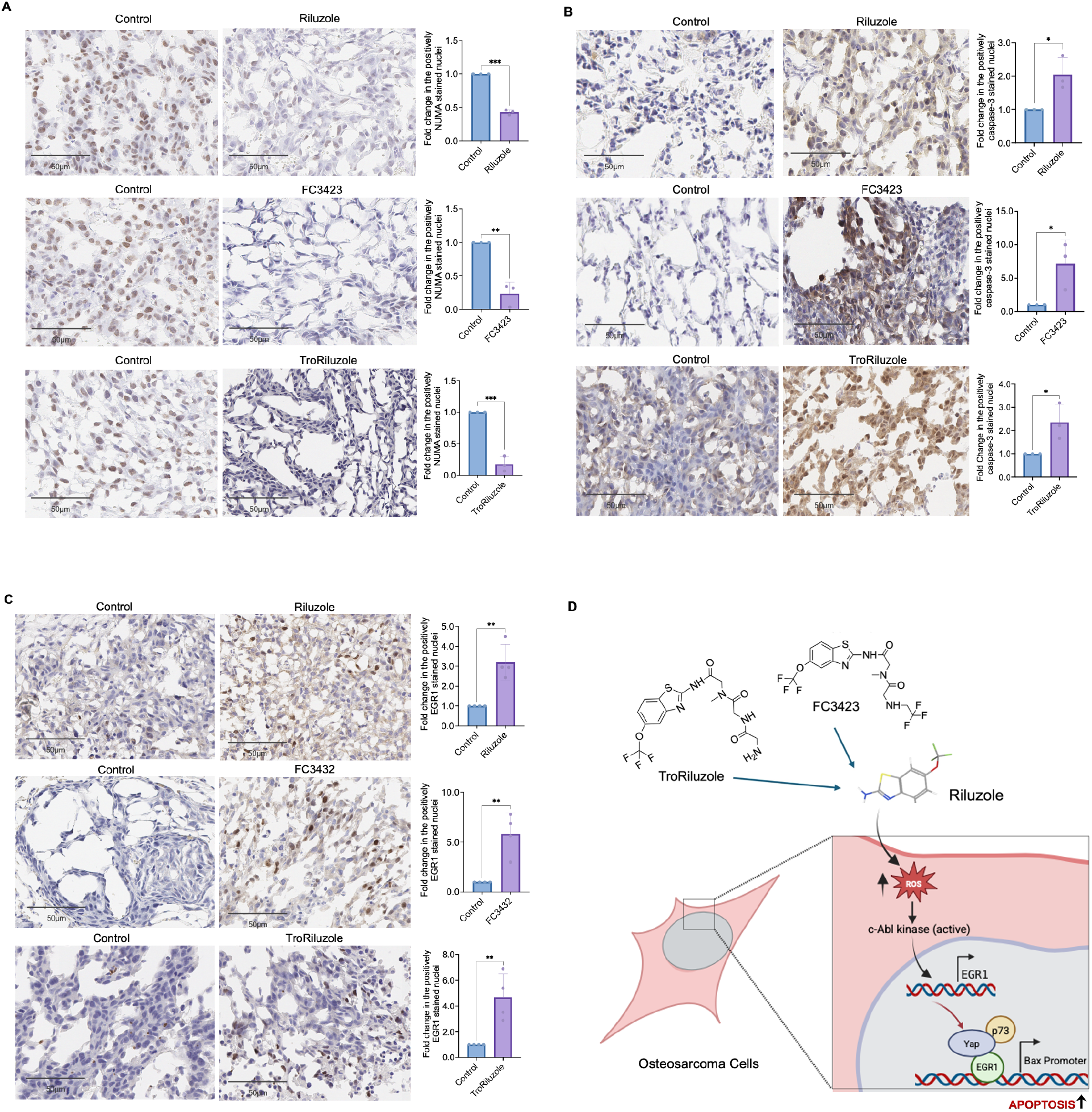
Riluzole enhances the expression of Egr-1 and Cleaved caspase 3 in the metastatic osteosarcoma mouse xenografts. Representative immunohistochemical images from all 4 groups: Control (left panels) and indicated drug treated (right panels, respectively). Scale bar = 50 μm. The bar graph (right panel) quantifies the number of positively stained, **(A)** NUMA, **(B)** cleaved caspase 3, **(C)** *EGR1*, cells in the treated groups compared to the control. Data are represented as mean SD of n=3, with statistical analysis performed using two-tailed Student’s t-test (*p<0.05, **p<0.005, ***p<0.0005). **(D)** Schematic diagram showing the mechanism of action of Riluzole in osteosarcoma cells: Riluzole induces the generation of reactive oxygen species (ROS) in OS cells through c-Abl kinase activity, leading to upregulation of the transcription factor *EGR1. EGR1* along with Yap/p73 activates the Bax gene expression promoting apoptosis.

## DISCUSSION

This study highlights the critical role of *EGR1* as a key mediator in facilitating the recruitment of the Yap/p73 complex to the Bax promoter and thereby driving apoptosis in Riluzole treated OS. Despite its known regulatory roles in cell growth, proliferation, and apoptosis across various cell types, the function of *EGR1* in OS remains poorly understood. Interestingly, *EGR1*, a transcription factor, exhibits dual roles in cancer biology, functioning either as an oncogene or a tumor suppressor depending on the cancer type (21, 22). For example, in prostate and gastric cancers, *EGR1* is markedly upregulated and acts as an oncogene, driving tumor progression. (21, 38). In contrast, *EGR1* activates pro-apoptotic genes leading to tumor cell apoptosis, hence serving as tumor suppressor in gliomas and pancreatic cancer (22, 39). Similarly, Yap, a transcriptional co-activator, can function both as oncogene and a tumor suppressor depending on its interacting partner (17, 31, 40). When Yap binds to TEAD proteins via its TEAD-binding domain (TBD), it acts as an oncogene (31). Conversely, when it interacts with p73 through its WW domain, specifically recognizing p73’s PPPY motif, Yap promotes tumor suppression (40, 41). Riluzole treatment shifts the pro-proliferative role of Yap to pro-apoptotic via phosphorylation of Yap at Y357 residue (17). Recent research also shows that Yap’s WW domain specifically binds to the PPxY motif of *EGR1*, which subsequently binds to the Bax promoter to induce apoptosis (24).

We initially concentrated on a panel of YAP-associated genes known to be critical regulators of apoptosis or proliferation. These included the pro-apoptotic genes *PUMA, PIG3*, and *PTEN*, as well as the pro-proliferative markers *CTGF* and *CYR61* (28-31, 42, 43). Riluzole treatment consistently led to a time-dependent increase in the pro-apoptotic genes and a corresponding decrease in proliferative markers. For example, in U2OS cells, both *PUMA* and *PIG3* were upregulated following Riluzole treatment. In LM7 cells, only *PUMA* was upregulated, suggesting that *PUMA* can mediate apoptosis in osteosarcoma independently of p53. *PUMA* and *PIG3* are pro-apoptotic genes regulated by p53, though they can also be activated independently of p53 (29, 44). Importantly, evidence from multiple studies confirms that p73, a p53 family member activated by Riluzole, directly binds to the promoter regions of key pro-apoptotic genes such as *PUMA* and *PIG3* (17, 44). For instance, p73 binding to the *PUMA* promoter has been demonstrated to transactivate *PUMA* expression in response to DNA damage, and similar binding has been reported for *PIG3*, emphasizing the role of p73 in triggering apoptosis even when p53 is compromised (44).

In contrast, *PTEN* was significantly upregulated in both cell lines (at 4 hours in U2OS and 6 hours in LM7), indicating its role in suppressing survival pathways. Indeed, it has been shown that *PTEN* is a direct target of *EGR1*, a transcription factor that interacts with YAP and regulates tumor suppressor networks (42). Increased *PTEN* expression reduces PIP3 levels, thereby suppressing the survival pathway and promoting apoptosis (45, 46).

Canonical YAP targets *CTGF* and *CYR61*, which promote proliferation, showed no early changes post-treatment; however, after 18 hours of Riluzole exposure, both genes were markedly downregulated in LM7 cells. This delayed suppression suggests diminished YAP-driven proliferative signaling. Overall, our data support a model where Riluzole promotes apoptosis by enhancing YAP and p73 activity, leading to the upregulation of pro-apoptotic genes such as *PUMA, PIG3*, and *PTEN*. Early *EGR1* induction may further amplify this response by acting as a transcriptional intermediary while repressing proliferation-associated genes like *CTGF* and *CYR61*. These findings identify *EGR1* as a tumor suppressor in osteosarcoma, with Riluzole-mediated upregulation driving apoptotic cell death.

We show that *EGR1* is downregulated in OS and that restoring its expression induces apoptosis, supporting its tumor-suppressive role. Riluzole significantly increased *EGR1* expression at both mRNA and protein levels across OS cell lines and two PDX lines, except in 143B cells. Our data indicate that Riluzole does not elevate ROS levels in 143B cells, in contrast to other OS lines (34). Notably, 143B cells harbor a KRAS mutation and are derived from HOS cells (47), the KARS mutation and lack of ROS may explain the absence of *EGR1* induction. This highlights ROS as a key upstream regulator of *EGR1*, further supported by NAC experiments where ROS scavenging attenuated *EGR1* upregulation.

Our prior study showed that Riluzole-induced ROS activates c-Abl kinase, facilitating YAP/p73 recruitment to the Bax promoter (17). A similar ROS-c-Abl mechanism has been reported to upregulate *EGR1* in U2OS cells (33). Here, we show Riluzole-induced ROS also increases *EGR1*, which binds to the Bax promoter and facilitates YAP/p73 recruitment, driving apoptosis. Interestingly, Bax expression rises only after *EGR1* begins to decline, suggesting a temporal cascade initiated by *EGR1*. Given the PPxY motif of *EGR1*, which can interact with YAP’s WW domain (24), we propose that *EGR1* helps stabilize the YAP/p73 complex at the Bax promoter, revealing a novel cooperative role in pro-apoptotic gene regulation.

Consistently, our *in vivo* metastatic OS mouse model validated these *in vitro* findings, showing Riluzole-induced *EGR1* upregulation and associated apoptosis. Previous studies reported that chemotherapies such as MAP and Etoposide induce *EGR1* and promote apoptosis (20). Our data show Riluzole and its prodrugs similarly increase *EGR1* in various OS models, positioning *EGR1* as a promising marker of drug efficacy. While current results are compelling, validation in patient tumor samples before and after Riluzole treatment is needed to confirm *EGR1*’s potential as a therapeutic biomarker.

## CONCLUSION

Our findings identify *EGR1* as a critical mediator of apoptosis in osteosarcoma and reveal that Riluzole-induced ROS enhances *EGR1* expression, promoting cell death via the Yap/p73-Bax axis. This study not only advances our understanding of *EGR1*’s tumor-suppressive role in osteosarcoma but also highlights its potential as a therapeutic marker. Further investigation into the upstream regulators and downstream effectors of *EGR1* may open new avenues for targeted therapies in osteosarcoma treatment.

## MATERIALS AND METHODS

### Cell lines and cell culture

Commercially available human OS cell lines (hFOB, SaOS-2, U2OS, MG63, HOS, 143B) from ATCC were employed as described previously (34). Cells were cultured in appropriate media, as previously described (34), supplemented with 10% FBS and 1% P/S, maintained at 37^°^C and 5% CO2. LM7 cells were provided by Dr. Eugenie Kleinerman, MD Anderson Cancer Center in Houston, Texas (48). MG63.3 cells were provided by Dr. Le Blanc from NCI/NIH. PDXs were from Dr. V.K Rajasekhar and Dr. J.H Healey, Memorial Sloan Kettering Cancer Center New York.

### Drug Treatments

For *in vitro* studies, all the cell lines were treated with 100 µM Riluzole for the *in vitro* experiments. For the antioxidant rescue experiments, 5mM of N-acetyl Cysteine (NAC) was used. For *in vivo* studies, we used Riluzole(7.5 mg/kg p.o QD), FC3423(4.3 mg/kg body p.o QD), TroRiluzole(4.1 mg/kg p.o QD). We received the pro-drugs FC3423 and TroRiluzole from Dr. Jeffery C. Pelletier and Dr. Allen B. Reitz at the Fox Chase Chemical Diversity Center, PA, USA (36).

### Cytotoxicity Assay

For MTT assay, 40,000 cells/mL were plated in a 24-well plate. The next day, cells were treated with various concentrations (0 µM, 5 µM, 10 µM, 25 µM, 50 µM, 100 µM) of Riluzole, TroRiluzole and FC3423 and the plate was then incubated for 48 hours at 37C. Followed by the addition of 100 µL of MTT reagent with a (5mg/mL) was added to each well and incubated for additional 2 hours at 37 °C. Then, the media was discarded, and the precipitate was dissolved in 1 mL of DMSO in each well. The absorbance for the plate was read using a microplate reader (Molecular Devices SpectraMax i3) at 520nm.

### Western Blot

The cell lysates were prepared using RIPA buffer and Protein was quantified before loading onto the gel. For transfer, PVDF membrane 0.45 µM was used. The blot was blocked with 5% non-fat dry milk in PBST buffer and incubated with primary antibodies *EGR1* (SCBT #sc-515830), Bax (Proteintech #50599-2-Ig) and GAPDH (Cell signaling #5174) in 5% BSA solution in PBST buffer, overnight at 4°C. The next day, the blot was washed three times with PBST buffer and incubated with secondary antibody in PBST buffer for 1 hour. ECL western blotting substrate was used for protein band detection. The blot was imaged using LICOR imager and the bands were quantified using LICOR Image Studio software.

### Quantitative Polymerase Chain Reaction (qPCR)

The cells were harvested using RLT lysis buffer (DTT freshly added) and RNA was purified using the RNeasy kit (Qiagen #74134). Briefly, the RNA was purified using the kit and then 1 µg of RNA was reverse transcribed to cDNA using Quantitech Reverse Transcription kit (Qiagen #205311). The cDNA was then diluted 10 times and then used for performing qPCR using SybrGreen Master Mix according to the manufacturer’s protocol.

### Chromatin Immunoprecipitation-qPCR

ChIP-qPCR was performed as previously described (17). Briefly, cells were treated with Riluzole for 0, 1, 2, 4 and 6 hours. At the end of each treatment, cells were fixed, washed, scrapped, collected in RIPA buffer and subjected to sonication. Prior to immunoprecipitation, 10% of the sonicated sample was set aside as input. Immunoprecipitation was performed overnight using A/G beads with antibodies for *EGR1* (Cell signaling #4153) at 1:50, YAP (Cell signaling #14074) at 1:50 dilution, p73 (Abcam #215038) at 0.544 mg/mL and pol II (Cell signaling #14958) at 1:50 dilution. The samples with only beads were used as control reference sample. Bax-promoter primers used were: Bax-promoter R: 5’-AGCTGCCACTCGGAAAAAGA-3’ and Bax-promoter F: 5’-AGGATGCGTCCACCAAGAAG-3’.

### Animal Study

The study was performed in accordance with the approved protocol by the Weill Cornell Medicine – MSKCC IACUC committee (IAUCUC protocol number: 2019-0049). We injected LM7-eGFP-ffLuciferase cells in PBS into the lateral tail veins of 5-week-old NOD.Cg-*Prkdc*^*scid*^ *Il2rg*^*tm1Wjl*^/SzJ female mice. On day 7, after the tumors have developed in the lungs, mice were randomly placed in 4 groups.

Group1(G1) had control mice, Group2(G2) mice received Riluzole, Group3(G3) received FC-3423 and Group4(G4) received TroRiluzole treatment through oral gavage. Mice were treated with PBS or Riluzole (7.5 mg/kg p.o QD), FC3423 (4.3 mg/kg body p.o QD), TroRiluzole (4.1 mg/kg p.o QD). Animals were given the drug daily by oral gavage and tumors were monitored on a weekly basis. The bioluminescence data were recorded for both ventral and dorsal flux for each mice every week. All experiments were performed in accordance with the approved IACUC protocol and other relevant guidelines and regulations. The maximal tumor size/burden was below the limits permitted by their IACUC ethics committee. The total flux (dorsal & ventral) was plotted for all 4 weeks and two-way ANOVA test was performed with Bonferroni’s post-hoc analysis to calculate the significant difference.

### Immunohistochemistry (IHC)

Chromogenic immunohistochemistry was performed on a Ventana Medical Systems Discovery Ultra platform using Ventana reagents and detection kits unless otherwise specified. Unconjugated polyclonal rabbit anti-human Nuclear Mitotic Apparatus Protein (NUMA) (Abcam Cat# ab97585 Lot# GR268490-46, RRID: AB_10680001), unconjugated rabbit anti-human Early Growth Response Protein-1 (*EGR1*) clone 15F7 (Cell Signaling Cat# 4153, Lot# 5, RRID: AB_2097038) and unconjugated rabbit anti-human and mouse Cleaved Caspase-3 (CASP) clone D3E9 (Cell Signaling Cat# 9579, Lot# 1, RRID: AB_10897512) antibody used for labeling. All samples were frozen OCT embedded and stored -80°C. Samples were sectioned at 5-microns onto Superfrost Plus (Fisher Scientific Cat# 1255015) or Superfrost Plus Gold (Fisher Scientific Cat# 1518848) slides and stored at -80°C prior to use. For immunohistochemistry, slides were removed from the freezer and allowed to dry overnight at room temperature. Samples were fixed in 10% Neutral Buffered Formalin (Fisher Scientific Cat# SF100-4) for 60 minutes at room temperature followed by rinsing in distilled water for 15 minutes. Slides were then incubated in Ventana Medical Systems Reaction Buffer (Ventana Medical Systems Cat# 950-300) for 30 minutes and subsequently loaded on to the instrument. Slides were treated with 3% hydrogen peroxide for 4 minutes to quench endogenous peroxidase. NUMA was diluted 1:7000 in Dulbecco’s PBS (Thermo-Fischer Cat# J67670) and incubated for 12 hours at room temperature. *EGR1* and CASP were diluted 1:100 in Cell Signaling Antibody Diluent (Cell Signaling Cat# 8112) and incubated for 3 hours at room temperature. NUMA, *EGR1* and CAS were detected with goat anti-rabbit Horseradish Peroxidase conjugated multimer (Ventana Cat# 760-4311) incubated for 8 minutes. Enzyme conjugated secondaries were detected with ChromoMap DAB detection (Cat# 760-159). Slides were washed in distilled water, counterstained with hematoxylin, dehydrated and mounted with permanent media. Negative controls consisted of primary antibody substituted with antibody diluent. All slides were scanned on an Aperio (Leica) AT2, captured at 40x and visualized on eSlideManager software (Version 12.5.0.6145). QuPath software was used for analyzing the IHC images.

## Statistical Analysis

The overall statistical analysis was performed using two-tailed Student’s *t-test*, one-way ANOVA with Dunnett’s post-hoc analysis and two-way ANOVA with Bonferroni post-hoc analysis.

## DECLARATIONS

### Ethics approval and consent to participate

Each patient signed a written consent preoperatively and the samples were collected using institutional review board–approved protocols (06-107 and 17-067).

## ACKNOWLEDGEMENTS

We thank Dr. Luis Chiriboga and his team for performing IHC at The NYULH Center for Biospecimen Research and Development, Histology and Immunohistochemistry Laboratory (RRID:SCR_018304) supported in part by the Laura and Isaac Perlmutter Cancer Center Support Grant; NIH/NCI P30CA016087. We thank Dr. Elisa De Stanchina at the anti-tumor assessment core facility at MSKCC for helping with the animal study. We would also like to thank Dr. Jeffery C. Pelletier and Dr. Allen B. Reitz for generously providing us with Riluzole pro-drugs FC3423 and TroRiluzole (36).

## REFERENCES

1. Sadykova LR, Ntekim AI, Muyangwa-Semenova M, Rutland CS, Jeyapalan JN, Blatt N, et al. Epidemiology and Risk Factors of Osteosarcoma. Cancer Invest. 2020;38(5):259–69.

2. Picci P. Osteosarcoma (osteogenic sarcoma). Orphanet J Rare Dis. 2007;2:6.

3. Savage SA, Mirabello L. Using epidemiology and genomics to understand osteosarcoma etiology. Sarcoma. 2011;2011:548151.

4. Meltzer PS, Helman LJ. New Horizons in the Treatment of Osteosarcoma. N Engl J Med. 2021;385(22):2066–76.

5. Misaghi A, Goldin A, Awad M, Kulidjian AA. Osteosarcoma: a comprehensive review. SICOT J. 2018;4:12.

6. Khan AJ, LaCava S, Mehta M, Schiff D, Thandoni A, Jhawar S, et al. The glutamate release inhibitor riluzole increases DNA damage and enhances cytotoxicity in human glioma cells, in vitro and in vivo. Oncotarget. 2019;10(29):2824–34.

7. Blyufer A, Lhamo S, Tam C, Tariq I, Thavornwatanayong T, Mahajan SS. Riluzole: A neuroprotective drug with potential as a novel anti-cancer agent (Review). Int J Oncol. 2021;59(5).

8. Saitoh Y, Takahashi Y. Riluzole for the treatment of amyotrophic lateral sclerosis. Neurodegener Dis Manag. 2020;10(6):343–55.

9. Crupi R, Impellizzeri D, Cuzzocrea S. Role of Metabotropic Glutamate Receptors in Neurological Disorders. Front Mol Neurosci. 2019;12:20.

10. Cowan RW, Seidlitz EP, Singh G. Glutamate signaling in healthy and diseased bone. Front Endocrinol (Lausanne). 2012;3:89.

11. Yang W, Maolin H, Jinmin Z, Zhe W. High expression of metabotropic glutamate receptor 4: correlation with clinicopathologic characteristics and prognosis of osteosarcoma. J Cancer Res Clin Oncol. 2014;140(3):419–26.

12. Robert SM, Sontheimer H. Glutamate transporters in the biology of malignant gliomas. Cell Mol Life Sci. 2014;71(10):1839–54.

13. Namkoong J, Shin SS, Lee HJ, Marin YE, Wall BA, Goydos JS, et al. Metabotropic glutamate receptor 1 and glutamate signaling in human melanoma. Cancer Res. 2007;67(5):2298–305.

14. Le MN, Chan JL, Rosenberg SA, Nabatian AS, Merrigan KT, Cohen-Solal KA, et al. The glutamate release inhibitor Riluzole decreases migration, invasion, and proliferation of melanoma cells. J Invest Dermatol. 2010;130(9):2240–9.

15. Genever PG, Skerry TM. Regulation of spontaneous glutamate release activity in osteoblastic cells and its role in differentiation and survival: evidence for intrinsic glutamatergic signaling in bone. FASEB J. 2001;15(9):1586–8.

16. Liao S, Ruiz Y, Gulzar H, Yelskaya Z, Ait Taouit L, Houssou M, et al. Osteosarcoma cell proliferation and survival requires mGluR5 receptor activity and is blocked by Riluzole. PLoS One. 2017;12(2):e0171256.

17. Raghubir M, Azeem SM, Hasnat R, Rahman CN, Wong L, Yan S, et al. Riluzole-induced apoptosis in osteosarcoma is mediated through Yes-associated protein upon phosphorylation by c-Abl Kinase. Sci Rep. 2021;11(1):20974.

18. Havis E, Duprez D. EGR1 Transcription Factor is a Multifaceted Regulator of Matrix Production in Tendons and Other Connective Tissues. Int J Mol Sci. 2020;21(5).

19. Gregg J, Fraizer G. Transcriptional Regulation of EGR1 by EGF and the ERK Signaling Pathway in Prostate Cancer Cells. Genes Cancer. 2011;2(9):900–9.

20. Matsunoshita Y, Ijiri K, Ishidou Y, Nagano S, Yamamoto T, Nagao H, et al. Suppression of osteosarcoma cell invasion by chemotherapy is mediated by urokinase plasminogen activator activity via up-regulation of EGR1. PLoS One. 2011;6(1):e16234.

21. Virolle T, Krones-Herzig A, Baron V, De Gregorio G, Adamson ED, Mercola D. Egr1 promotes growth and survival of prostate cancer cells. Identification of novel Egr1 target genes. J Biol Chem. 2003;278(14):11802–10.

22. Wang C, Husain K, Zhang A, Centeno BA, Chen DT, Tong Z, et al. EGR-1/Bax pathway plays a role in vitamin E delta-tocotrienol-induced apoptosis in pancreatic cancer cells. J Nutr Biochem. 2015;26(8):797–807.

23. Saha SK, Islam SMR, Saha T, Nishat A, Biswas PK, Gil M, et al. Prognostic role of EGR1 in breast cancer: a systematic review. BMB Rep. 2021;54(10):497–504.

24. Zagurovskaya M, Shareef MM, Das A, Reeves A, Gupta S, Sudol M, et al. EGR-1 forms a complex with YAP-1 and upregulates Bax expression in irradiated prostate carcinoma cells. Oncogene. 2009;28(8):1121–31.

25. Windischhofer W, Huber E, Rossmann C, Semlitsch M, Kitz K, Rauh A, et al. LPA-induced suppression of periostin in human osteosarcoma cells is mediated by the LPA(1)/Egr-1 axis. Biochimie. 2012;94(9):1997–2005.

26. Jung SN, Oh C, Chang JW, Liu L, Lim MA, Jin YL, et al. EGR1/GADD45alpha Activation by ROS of Non-Thermal Plasma Mediates Cell Death in Thyroid Carcinoma. Cancers (Basel). 2021;13(2).

27. Chen Z, Wu FF, Li J, Dong JB, He HY, Li XF, et al. Investigating the synergy of Shikonin and Valproic acid in inducing apoptosis of osteosarcoma cells via ROS-mediated EGR1 expression. Phytomedicine. 2024;126:155459.

28. Baron V, Adamson ED, Calogero A, Ragona G, Mercola D. The transcription factor Egr1 is a direct regulator of multiple tumor suppressors including TGFbeta1, PTEN, p53, and fibronectin. Cancer Gene Ther. 2006;13(2):115–24.

29. Jeffers JR, Parganas E, Lee Y, Yang C, Wang J, Brennan J, et al. Puma is an essential mediator of p53-dependent and -independent apoptotic pathways. Cancer Cell. 2003;4(4):321–8.

30. Zhang C, Wei W, Tu S, Liang B, Li C, Li Y, et al. Upregulation of CYR61 by TGF-beta and YAP signaling exerts a counter-suppression of hepatocellular carcinoma. J Biol Chem. 2024;300(4):107208.

31. Zhao B, Ye X, Yu J, Li L, Li W, Li S, et al. TEAD mediates YAP-dependent gene induction and growth control. Genes Dev. 2008;22(14):1962–71.

32. Shome D, von Woedtke T, Riedel K, Masur K. The HIPPO Transducer YAP and Its Targets CTGF and Cyr61 Drive a Paracrine Signalling in Cold Atmospheric Plasma-Mediated Wound Healing. Oxid Med Cell Longev. 2020;2020:4910280.

33. Stuart JR, Kawai H, Tsai KK, Chuang EY, Yuan ZM. c-Abl regulates early growth response protein (EGR1) in response to oxidative stress. Oncogene. 2005;24(55):8085–92.

34. Jung O, Rajasekhar VK, Azeem SM, ChandThakuri S, Norton B, Healey JH, et al. Repurposing riluzole as an anti-osteosarcoma agent. Frontiers in Oncology. 2025;Volume 15 - 2025.

35. Ajroud-Driss S, Saeed M, Khan H, Siddique N, Hung WY, Sufit R, et al. Riluzole metabolism and CYP1A1/2 polymorphisms in patients with ALS. Amyotroph Lateral Scler. 2007;8(5):305–9.

36. Pelletier JC, Chen S, Bian H, Shah R, Smith GR, Wrobel JE, et al. Dipeptide Prodrugs of the Glutamate Modulator Riluzole. ACS Med Chem Lett. 2018;9(7):752–6.

37. Raghubir M, Rahman CN, Fang J, Matsui H, Mahajan SS. Osteosarcoma growth suppression by riluzole delivery via iron oxide nanocage in nude mice. Oncol Rep. 2020;43(1):169–76.

38. Li L, Ameri AH, Wang S, Jansson KH, Casey OM, Yang Q, et al. EGR1 regulates angiogenic and osteoclastogenic factors in prostate cancer and promotes metastasis. Oncogene. 2019;38(35):6241–55.

39. Calogero A, Arcella A, De Gregorio G, Porcellini A, Mercola D, Liu C, et al. The early growth response gene EGR-1 behaves as a suppressor gene that is down-regulated independent of ARF/Mdm2 but not p53 alterations in fresh human gliomas. Clin Cancer Res. 2001;7(9):2788–96.

40. Zhang H, Wu S, Xing D. YAP accelerates Abeta(25-35)-induced apoptosis through upregulation of Bax expression by interaction with p73. Apoptosis. 2011;16(8):808–21.

41. Strano S, Munarriz E, Rossi M, Castagnoli L, Shaul Y, Sacchi A, et al. Physical interaction with Yes-associated protein enhances p73 transcriptional activity. J Biol Chem. 2001;276(18):15164–73.

42. Virolle T, Adamson ED, Baron V, Birle D, Mercola D, Mustelin T, et al. The Egr-1 transcription factor directly activates PTEN during irradiation-induced signalling. Nat Cell Biol. 2001;3(12):1124–8.

43. Flatt PM, Polyak K, Tang LJ, Scatena CD, Westfall MD, Rubinstein LA, et al. p53-dependent expression of PIG3 during proliferation, genotoxic stress, and reversible growth arrest. Cancer Lett. 2000;156(1):63–72.

44. Klanrit P, Flinterman MB, Odell EW, Melino G, Killick R, Norris JS, et al. Specific isoforms of p73 control the induction of cell death induced by the viral proteins, E1A or apoptin. Cell Cycle. 2008;7(2):205–15.

45. Weng L, Brown J, Eng C. PTEN induces apoptosis and cell cycle arrest through phosphoinositol-3-kinase/Akt-dependent and -independent pathways. Hum Mol Genet. 2001;10(3):237–42.

46. Stambolic V, Suzuki A, de la Pompa JL, Brothers GM, Mirtsos C, Sasaki T, et al. Negative regulation of PKB/Akt-dependent cell survival by the tumor suppressor PTEN. Cell. 1998;95(1):29–39.

47. Hensler PJ, Annab LA, Barrett JC, Pereira-Smith OM. A gene involved in control of human cellular senescence on human chromosome 1q. Mol Cell Biol. 1994;14(4):2291–7.

48. Jia SF, Worth LL, Kleinerman ES. A nude mouse model of human osteosarcoma lung metastases for evaluating new therapeutic strategies. Clin Exp Metastasis. 1999;17(6):501–6.

